# Genomic selection performs as effectively as phenotypic selection for increasing seed yield in soybean

**DOI:** 10.1101/2022.07.12.499836

**Authors:** Nonoy B. Bandillo, Diego Jarquin, Luis G. Posadas, Aaron J. Lorenz, George L. Graef

## Abstract

Increasing the rate of genetic gain for seed yield remains the primary breeding objective in both public and private soybean breeding programs. Genomic selection (GS) has the potential to accelerate the rate of genetic gain for soybean seed yield. To date, limited studies have empirically validated accuracy of GS and compared to phenotypic selection (PS), and none has been done for soybean breeding. This study conducted the first empirical validation of GS for increasing seed yield using over 1,500 lines and over 7 years (2010-2016) of replicated experiments in the University of Nebraska soybean breeding program. The study was designed to capture the varying genetic relatedness of the training population to three validation sets: two large bi-parental populations (TBP-1 and TBP-2), and a large validation set comprised of 457 pre-selected advanced lines derived from 45 bi-parental populations in the variety development program (TMP). We found that prediction accuracy (0.54) from our validation experiments was competitive with what we obtained from a series of cross-validation experiments (0.64). Both GS and PS were more effective for increasing population mean performance with similar realized gain but significantly greater than random selection (RS). We found a selection advantage of GS over PS where higher genetic gain and identification of top-performing lines was maximized at higher selection stringency from 10 to 20% selected proportion. GS led to at least 2% increase in the mean genetic similarity vs. PS and RS, potentially causing a minimal loss of genetic diversity. We showed that loss of genetic variance in the GS set was presumably due to a significant shift on allelic frequencies towards the extremes. Across all loci, an average increase of 0.04 in allelic frequency in the GS set was observed after selection, which is about 5% higher than the base population when no selection was made. Overall, we demonstrate that GS performed as effectively as PS, and the implementation of GS in a public soybean breeding program should be warranted mainly for reducing breeding cycle time and lowering cost per unit gain.

## Introduction

Increasing the rate of genetic gain for seed yield remains the primary breeding objective in both public and private soybean breeding programs. Similar with virtually all major crops, the genetic architecture of soybean seed yield is highly complex, controlled by many loci of small effects, which complicates breeding and effective selection. A recent report of genetic gain for soybean seed yield showed ~29 kg ha^−1^ yr^−1^ (0.43 bu ha^−1^ yr^−1^) using a historic set of MG II-IV soybean varieties released from 1923 to 2008 (Rinker et al., 2014; Specht et al., 2014). The increment in genetic gain for soybean seed yield is lower than the reported genetic gain for corn seed yield, which generally ranges from ~63 to 75 kg ha^−1^ yr^−1^ (1.0 to 1.2 bu ha^−1^ yr^−1^) (Duvick., 1984, 2005a, 2005b; Bubeck et al., 2006; Smith et al., 2014), but both crops have an average gain in yield of about 0.8% per year. Similar to other staple crops, this rate of genetic gain is not sufficient to cope with the 2% yearly increase in the world population (Hickey et al 2016), which is further exacerbated by changing environmental conditions. The major challenge is how to continuously increase the rate of genetic gain of soybean seed yield to more than double relative to the current levels (Diers et al., 2018).

Historically, increase in genetic gain in soybean is mainly attained through selective breeding and with the integration of results from other disciplines in agricultural research, including agronomic technology and innovation, crop and soil management practices, and plant pathology, to mention a few (Specht et al., 2014). Soybean breeders also have increased the annual rate of genetic gain through use of off-season winter nurseries, which substantially reduces number of years needed to develop new soybean cultivars. The most common breeding methods that have been applied for advancement and/or selection in public soybean breeding programs are a combination of single-seed descent, pedigree, and bulked method, which sometimes involves visual selection and trait screening (e.g., SCN, iron deficiency chlorosis, etc.) as early as F_2_ or over successive generations. Phenotypic selection methods for seed yield and other important agronomic traits involve the use classical statistical design and research plots in single or multi-environment (location-year) trials that are representative of the target population of environments (TPE) for the area of adaptation, mostly determined by latitude. With current advances in soybean genetics and genomics, other potentially faster breeding methods have been developed (Song et al 2013; 2015). Marker-assisted selection (MAS) for simple Mendelian and oligogenic traits such as maturity (Bandillo et al., 2017; Cao et al., 2017; Lu et al., 2017; Fang et al., 2021), descriptive traits (Bandillo et al., 2017), and disease resistance (e.g., SCN) has recently been used for genetic improvement in soybean. However, its overall impact to increase the rate of genetic gain for soybean seed yield has been limited.

Genomic selection (GS) takes advantage of high-density genomic data and holds a promise of identifying and selecting candidates with the best genotypic value for complex traits such as seed yield. GS has become an attractive option as a selection-decision tool in plant breeding programs as genotyping costs have significantly declined to the extent that the per-individual cost of sequencing is less expensive than the per-sample cost of multi-environment testing (Ertiro et al., 2015; Crossa et al., 2017; Bernardo, 2020). With the plummeting cost of sequencing coupled with continued improvement on prediction models and high-performance computing, assessing prediction accuracy through strategic cross-validation schemes on available germplasm or breeding materials has now become a routine in crop species (Bernardo, 2016; Crossa et al., 2017). To date, however, limited studies have empirically validated the accuracy of GS relative to PS, and only a few has directly implemented GS for applied plant breeding efforts (Massman et al., 2013; Asoro et al., 2013; Combs and Bernardo, 2013; Beyene et al, 2015; Rutkoski et al., 2015). Most of these empirical studies have showed that the accuracy of GS was no better than that of PS, and the observed genetic gain was consequently non-significant between GS and PS (Sallam and Smith, 2016; Rutkoski et al., 2015; Heffner et al., 2011b). On the contrary, Massman et al. (2013) showed that up to 50% greater genetic gain can be achieved from GS when compared with marker-assisted recurrent selection, whereas Asoro et al. (2013) found that selection using molecular markers and phenotype jointly was significantly more effective than selection based on pedigree and phenotype. More empirical data are needed to validate selection advantages of GS over PS, and such empirical type of study has been lacking in soybean breeding.

In this study, we built upon from the work of Jarquin et al. (2014) and conducted a follow-up study to empirically validate the accuracy of GS for increasing seed yield in the University of Nebraska soybean breeding program using over 1,500 lines and over 7 years (2010-2016) of replicated experiments. Jarquin et al. (2014) estimated a maximum prediction ability for seed yield of 0.64 using extensive cross-validation experiments with the TP used in this study. To verify whether this level of prediction accuracy would be realized in practice, three validation sets were used including two independent bi-parental populations (hereafter, TBP-1 and TBP-2), and an independent validation set (TMP) comprised of pre-selected advanced lines derived from multiple bi-parental populations. We compared GS, PS and random selection (RS) methods for improving the population mean and genetic gain for soybean seed yield. We then assessed the impact of different selection methods on the level of genetic similarity and changes on allele frequency spectrum.

## Materials and Methods

### Plant materials

Multiple sets of breeding populations developed by the University of Nebraska-Lincoln soybean breeding program were utilized for this study (**Figure 1**, **Table S1).** First, the training population, hereafter TP, was used to train prediction models for target traits of interest (Jarquin et al., 2014). Second, a validation set was extracted from the program, which comprised pre-selected, advanced lines derived from multiple bi-parental populations, hereafter TMP. Third, two additional independent bi-parental populations (hereafter, TBP-1 and TBP-2) were also developed for the validation experiment. TBP-1 represented a population less closely related to the TP, and TBP-2 represented a population with greater genetic similarity to the TP among the range of populations in the program. The selected breeding materials from three validation sets (TMP, TBP-1, and TBP-2) were used to compare genomic selection (GS), phenotypic selection (PS), and random selection (RS). These breeding materials were evaluated in multiple experiments across seven years between 2010-2016 (**Fig. 1, Table S1**), which are described in more detail below.

**Figure 1.**
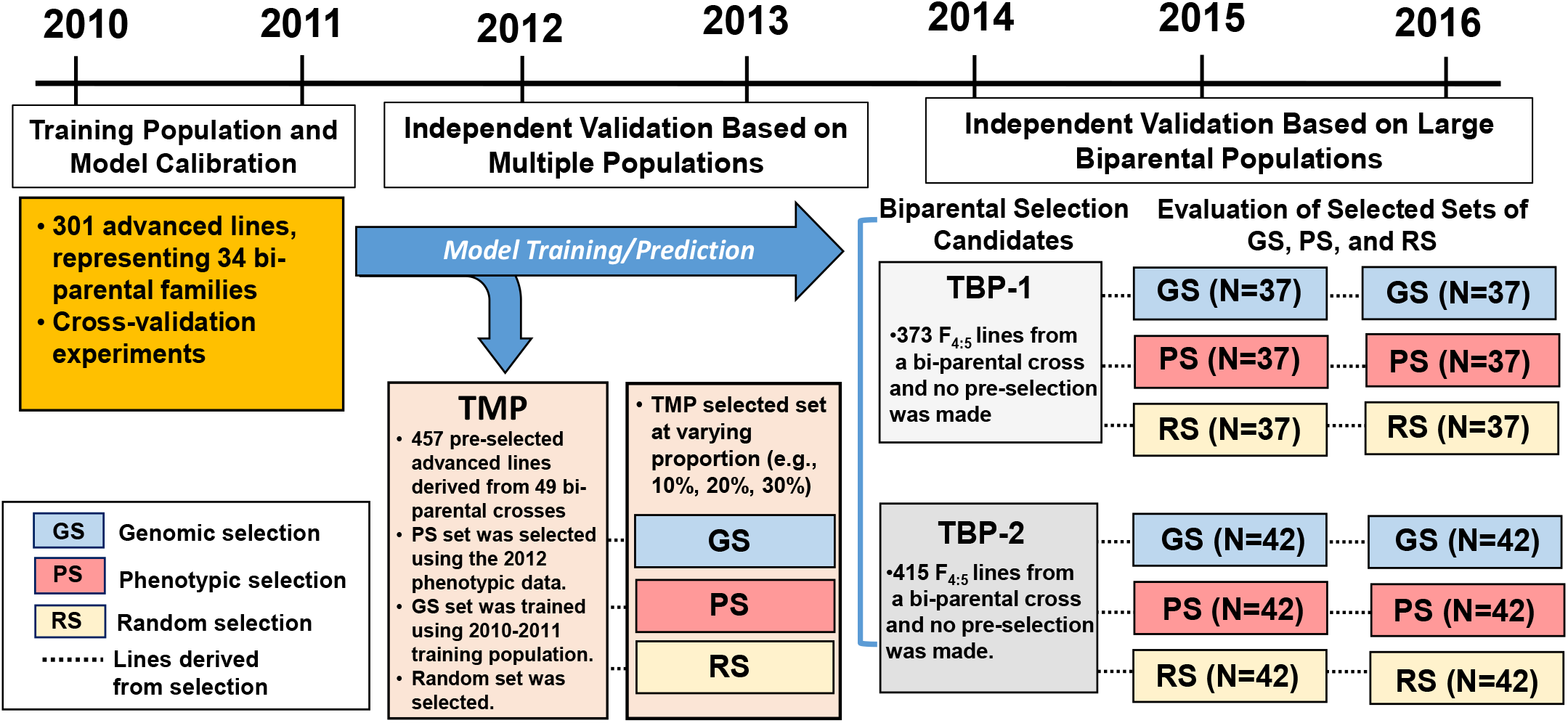
Study design and schematic overview of the project to empirically validate prediction accuracy and compare genomic selection (GS) to phenotypic selection (PS) and random selection (RS) in the University of Nebraska Soybean Breeding Program.

The first experiment was the extraction and evaluation of the TP in 2010 and 2011 to generate the data for training and optimization of prediction models. The TP used to generate prediction models was comprised of 301 experimental breeding lines described in Jarquin et al. (2014). Briefly, 272 lines were in the F_5:8_ generation and 26 in the F_4:7_ generation. Soybean lines belonged to maturity groups I (N = 64), II (N = 213), and III (N = 24) and represent 34 bi-parental families ranging in size from one to 28 lines per family. In that study, different models and cross-validation schemes were used to evaluate the accuracy of genomic predictions.

The second experiment was the extraction and evaluation of TMP in 2012 and 2013 to generate phenotypic data and empirically validate prediction accuracy. TMP consisted of pre-selected 457 F_4:5_ lines based on seed yield and other-agronomic characteristics (e.g., maturity, lodging, plant height), which were mostly derived from a set of crosses among breeding lines from the TP. The pre-selected set was comprised of breeding lines that were advanced to the second stage of testing in the program based on phenotypic data for yield. There were forty-nine unique lines represented as parents in these lines, representing 45 different biparental cross combinations. All the breeding populations included in TMP were advanced using single-seed descent method without selection up to F_4_, at which point lines were derived.

The third experiment was the development and evaluation of two additional independent bi-parental populations (TBP-1 and TBP-2). TBP-1 was derived from a cross between U09-105007 and IA4005, lines part of the TP set, and comprised of 373 F_4:5_ lines; and (2)TBP-2, consisted of 415 F_4:5_ lines and was derived from a cross between U09-215057 and LD07-3419. All breeding lines were advanced through single seed descent without selection up to F_4_, at which point lines were derived. In 2014, all F_4:5_ lines from the two TBP sets were included in preliminary yield trial (PYT) for evaluation of seed yield and other agronomic traits.

### Phenotypic evaluation of TP, TMP and TBPs

The number of breeding lines, years of evaluation, experimental design and number of environments (location X year) are displayed in **Table S1**. Phenotypic evaluation for the TP was previously described by Jarquin et al. (2014). The TP was evaluated in 2010 and 2011. Briefly, an augmented incomplete block design with two replications was used. At least three common checks were included in each block to correct for spatial field variability. Blocks generally consisted of at least 27 experimental entries. Lines belonging to maturity groups I and II were evaluated at the Nebraska locations Beemer, Phillips, Cotesfield, and Mead. Lines belonging to maturity group III were evaluated at the Nebraska locations Phillips, Mead, Lincoln, and Clay Center. Grain yield was measured at all locations and recorded as machine harvestable grain yield adjusted to 13% moisture.

TMP, TBP-1 and TBP-2 were evaluated following the same experimental design for the TP as described above, but differed only in years of evaluation and number of environments. TMP was evaluated in 2012 and 2013, whereas both TBP-1 and TBP-2 were evaluated in the same experiment in 2014 (Figure 1, Table S1). Test lines and checks were grown in four-row plots (0.76 m apart, 2.9 m long) with planting density of 26 seeds per square meter. Blocks consisted of 25-35 experimental entries and three check cultivars. Lines and common checks were assigned randomly within each incomplete block. Grain yield was measured at all locations and recorded as machine harvestable grain yield adjusted to 13% moisture.

### Genotyping and genetic relatedness

The DNA extraction protocol used for this study was previously described (Jarquin et al., 2014). Briefly, leaf discs were collected from 12 random plants of each soybean line at approximately the V6 growth stage. DNA isolation was performed using the Qiagen DNeasy Plant 96 kit following their recommended extraction protocol. All extracted DNA were sent to the Institute of Genomic Diversity at Cornell University for genotyping-by-sequencing (GBS) following the protocol described by Elshire et al. (2011).

The GBS pipeline implemented in TASSEL version 4.3.2 (Glaubitz et al., 2014) was used to discover SNP in all the population sets. We adopted the SNP calling parameters described by Jarquin et al. (2014). Briefly, tag counts were generated and merged from fastq files. Tag counts were then indexed and aligned to the reference genome Gmax_109_softmasked.fa.gz, which was downloaded from ftp://ftp.jgi-psf.org/pub/JGI_data/phytozome/v8.0/Gmax/assembly/ with BWA version 0.7.3a-r367. Scaffolds were ignored for SNP calling. All the analyses were performed through the Holland Computing Centre at the University of Nebraska-Lincoln.

Out 209,994 called SNP across TP and validation sets, we retained a substantial number of SNP that spanned across the genome after filtering based on percent missing values (PMV) and minor allele frequency (MAF) cut-off (**Figure S1**). We obtained a total of 20,670 common SNPs with PMV ≤ 80 % and MAF > 0.01 among TP and validation sets. Using these 20,670 common SNPs, we built our prediction models and analysis of population structure.

### Extraction of phenotypic, genomic, and random selection sets

Three validation sets (TMP, TBP-1 and TBP-2) were used to compare the effectiveness of PS, GS, and RS (**Table S1; Figure 1).** The top target-% performing individuals were selected by each method within each population set.

#### Phenotypic selection set

To form the PS selected set, we used the 2013 PYT data generated for TMP, and the 2014 PYT data for both the TBP sets. This mimics the timing of selection in practice where at least 10-20% selection intensity is used in a typical year based on the PYT data. Selected lines are then entered into replicated advanced yield trials. Ranking and selection of test entries were performed within each PS population (e.g., TMP, TBP-1, and TBP-2) based on calculated BLUPs following a linear mixed model below:

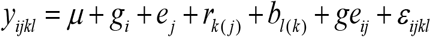

where *g_i_* represents the effect of the *i*^th^ genotype (i.e., soybean line), *e_j_*, represents the effect of the *j*^th^ location, *r_k(j)_* represents the effect of the *k*^th^ replicate nested in location *j*, *b_l(k)_* represents the effect of the *l*^th^ incomplete block nested within replicate *k, ge_ij_*, represent the effect of the *i*^th^ genotype tested in *j*^th^ location, and *ε_ijkl_* represents the residual term capturing the non-explained variability and it is assumed to be an independent and identically distributed (IID) random outcome following a normal density. The top target-% performing individuals was selected by each method within each population set.

#### Genomic selection set

To form the GS selected set, the RR-BLUP genomic prediction model was trained. A simpler method like RR-BLUP could be a good approximation to the optimal model, and Prediction accuracy values were generally similar among prediction models despite varying level of complexity (Jarquin et al., 2014; Yu et al., 2016). Unless otherwise indicated, all genomic predictions were performed using the RR-BLUP model based on the TP described by Jarquin et al. (2014). Using the shared SNPs between the TP and validation sets, the RR-BLUP model was trained using the equation

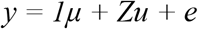

where y is a vector of BLUPs based on the 2010 and 2011 data generated for the TP (Jarquin et al., 2014); *μ* is an intercept vector; Z is an *n* × *p* incidence matrix containing the allelic states of the *p* marker loci (*z* = {-1, 0, 1}), where −1 represents the minor allele; u is the *p* × 1 vector of marker effects; and *e* is a *n* × 1 vector of residuals. Under RR-BLUP, *u* ~ MVN (0, I*σ*^2^_*u*_) where *σ*^2^_*u*_ is the variance of the common distribution of marker effects and was estimated using restricted maximum likelihood. The RR-BLUP model was implemented in the R package rrBLUP version 4.2 (R Development Core Team, 2012; Endelman, 2011). Prediction of the individuals were calculated independently for each TMP and TBP set using the equation

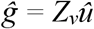

where 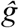 is a vector of genomic estimated breeding values (GEBV), *Z_v_* is the marker incidence matrix of TMP and TBP sets, and *û* is the vector of marker effect derived from training the RR-BLUP model with the TP set. The GS set was formed by selecting the top target-% lines within each TMP-GS and TBPs-GS set based on the ranked GEBVs for seed yield. Similar with PS and RS, the GS set had 37 selected lines for TBP-1, whereas 42 selected lines for TBP-2.

#### Random selection set

The RS set, which was used as control set, was formed by randomly sampling a similar proportion of 10% (unless otherwise specified) from all the individuals included in each of TMP and the two TBP sets. Each selected set by PS and RS consisted of the same number of lines as the GS sets; 46 selected individuals in TMP at 10% selection, 92 individuals in TMP at 20% selection and 138 individuals in TMP at 30% selection. TBP-1 had 37 selected lines while TBP-2 comprised of 42 selected lines.

### Phenotypic evaluation of phenotypic, genomic and random selected sets

The full set of TMP were evaluated in 2012 and 2013. That allowed us to explore the varying selected proportion for GS, PS and RS. In TBP-1 and TBP-2, only the top 10% selected lines by GS, PS and RS were evaluated in the same experiment in 2015 and 2016. All selected lines were evaluated in 8 environments of their corresponding maturity using an augmented incomplete block design with two replications (**Table S1**). Lines belonging to maturity groups I and II were evaluated at the Nebraska locations Beemer, Phillips, Cotesfield, and Mead. Lines belong to maturity group III were evaluated at the Nebraska locations Phillips, Mead, Lincoln, and Clay Center. Test lines and checks were grown in four-row plots (0.76 m apart, 2.9 m long) with planting density of 26 seeds per square meter. Blocks consisted of 25-35 experimental entries and three check cultivars. Lines and common checks were assigned randomly within each incomplete block. Grain yield was measured at all locations and recorded as machine harvestable grain yield adjusted to 13% moisture.

### Phenotypic data analysis and heritability

Different population sets were not evaluated in the same set of environments. The TP was evaluated in 2010 and 2011; TMP was evaluated in 2012-2013; whereas the TBP-1 and TBP-2 were evaluated in 2014-2016 (**Fig. 1; Table S1**). All experiments, however, were laid out following the same experimental design. Unless otherwise stated, all phenotypic data analysis was conducted using the same linear mixed model described below:

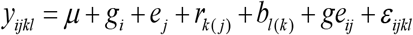

where *g_i_* represents the effect of the *i*^th^ genotype (i.e., soybean line), *e_j_* represents the effect of the *j*^th^ location, *r_k(j)_* represents the effect of the *k*^th^ replicate nested in location *j*, *b_l(k)_* represents the effect of the *l*^th^ incomplete block nested within replicate *k, ge_ij_* represent the effect of the *i*^th^ genotype tested in *j*^th^ location, and *ε_ijkl_* represents the residual assumed to be IID random outcomes from a normal density.

For the purpose of estimating heritability, we fit all factors as random effects. Variance components were estimated using the R-ASREML (Butler et al 2009?). The heritability on an entry-mean basis was estimated as

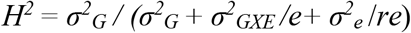

where *σ*^2^*G* is the genotypic variance, *σ^2^GXE* is the genotype-by-environment interaction, *σ^2^_e_* is the residual variance*, e* represents the number of environments and *r* is the number of replications within each environment.

### Evaluation of selection methods

Using selected sets from TMP and the two TBP sets, different metrics were used to compare the effectiveness of PS, GS, and RS, which are described in more detail below. We excluded the data when phenotypic selection was made as it would introduce biases toward PS. For example, TMP-PS was evaluated as part of the full TMP in 2012 for TMP and TBP-PS sets were evaluated as part of the full TBP sets in 2014 for the SC sets. We excluded 2012 and 2014 data in the evaluation of selected sets, and used only the 2013 data for TMP-PS, and the 2015 and 2016 data for the TBP-PS sets. The same principle was applied to RS sets.

1. **Mean population performance and ranking of high- and low-yielding genotypes.** Mean population and ranking of entries were performed within each population set. The calculated BLUE across years were used to get mean performance within each selection method. Entries within each population were ranked from most-to least-favorable individuals. Mean separation tests of selection methods were conducted using Tukey’s honestly significant difference (a=0.05).
2. **Realized gain from selections and relative efficiency.** Realized gains were calculated by subtracting the mean of selected sets (e.g., GS, PS, RS) from the mean of the full (All) set evaluated at year *t*, where *t* represents the year when both selected and all candidate lines were evaluated together as a full set. Percentage gain was calculated as realized gain divided by the mean of the full set. Paired two-tailed t-tests were used to test differences in gain between GS and PS, as well as differences between observed and expected gains. One-sided, one-sample t-test was used to test if genetic gains were significant. The relative efficiency of GS to PS (RE_GS:PS_) was calculated as the ratio of the response to GS at year *t* divided by the response to PS at year *t*.
3. **Genetic similarity among selected sets.** The simple matching coefficient (Sokal and Michener, 1958) was used to determine the genetic similarity based on 20,670 shared SNPs. The analysis was implemented in the R-package nomclust (Zulc and Rezankova, 2017).
4. **Distribution of allele frequency before and after selection.** We hypothesized that allele frequencies are shifted up to some degree after selection. To evaluate such hypothesis, we estimated and compared the distribution of allele frequency before (full set) and after selection (GS, PS, RS) in TMP, TBP-1 and TBP-2. Allele frequency before selection was estimated using all lines without selection, whereas allele frequency after selection was estimated using lines selected by each selection method. Mean allelic frequency across all SNPs was compared using Tukey’s honestly significant difference (a=0.05). A two-sample, two-sided Kolmogorov-Smirnov (KS) test at a=0.05 was used to compare cumulative allele frequency spectrum between the full set and selected sets.

#### Maximizing the prediction reliability

We evaluated the potential of using concept of reliability in the genomic best linear unbiased prediction (GBLUP; VanRaden 2008) model. Prediction reliability (VanRaden, 2008) was obtained for TMP, TBP-1, and TBP-2 using 20,670 shared SNP with the TP. The reliability criteria for each validation set (e.g., TMP, TBP-1, and TBP-2) were then calculated using the formula (Hayes et al., 2009; Yu et al., 2016) as follows:

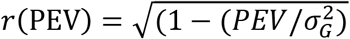

where PEV is the predicted error variance, and 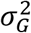 is the genetic variance.

## Results

### Heritability

Broad-sense heritability (*H^2^*) for seed yield in different population sets and environments are presented in **Table S2**. *H^2^* estimates for seed yield were not consistent among years but on general high values were observed in each population set. In general values ranged from 0.47 to 0.80. *H^2^* for seed yield in TP was 0.77, whereas in TMP *H^2^* ranged from 0.63 to 0.80. Similarly, high *H^2^* values were observed for TBP-1 and TBP-2 that ranged from 0.63 to 0.66.

### Prediction accuracy from empirical validation experiments

First, we looked at prediction accuracy for seed yield in TMP, which represents a good validation set as it was comprised of pre-selected lines closely related from the TP. Based on the top two principal components that captured 14% of total genetic variation, we did not observe any apparent population structure between the TP and TMP (**Figure S2**). Prediction accuracy in TMP ranged from 0.27 to 0.42 (**Table 1**) and, in most cases, was higher in 2012-2013 (0.42±0.03) than either 2012 (0.42±0.05) or 2013 alone (0.31±0.04) (**Table 1**).

**Table 1.** Prediction accuracy for seed yield in three full validation sets, and all the selected sets by GS, PS, and RS. Number of breeding lines (N) and years of evaluation are indicated in each set.

| Validation Set | N | Years of Evaluation | Training Population Set |  |  |
| --- | --- | --- | --- | --- | --- |
|  |  |  | TP 2010 | TP 2011 | TP 2010-2011 |
|  |  |  | ----- Prediction Accuracy (Standard Error) ----- |  |  |
| TMP (before selection) | 457 | 2012 | 0.32 (0.05) | 0.42 (0.05) | 0.39 (0.05) |
| TMP selected set |  | 2013 | 0.31 (0.04) | 0.28 (0.04) | 0.27 (0.04) |
| TMP selected set |  | 2012-2013 | 0.39 (0.04) | 0.42 (0.03) | 0.41 (0.04) |
| TBP-1 (before selection) | 373 | 2014 | 0.54 (0.05) | 0.16 (0.05) | 0.43 (0.05) |
| TBP-1 selected set | 99† | 2015-2016 | 0.32 (0.10) | 0.25 (0.10) | 0.27 (0.10) |
| TBP-2 (before selection) | 415 | 2014 | 0.04 (0.06) | 0.11 (0.01) | 0.13 (0.05) |
| TBP-2 selected set | 115† | 2015-2016 | 0.29 (0.09) | 0.17 (0.09) | 0.23 (0.09) |

†If a breeding line was selected more than once by at least one selection method, genotypic duplicates were dropped but retained one in a subsequent evaluation trial.

We then looked at prediction accuracy in the other two independent bi-parental populations, TBP-1 and TBP-2. As expected, breeding lines from the same cross clustered together (**Figure S2)**. TBP-1 was more genetically related to TP and TMP, whereas TBP-2 was less related to TP and TMP (**Figure S2)**. Prediction accuracy before selection for TBP-1 ranged from 0.16 to 0.54, whereas prediction accuracy for TBP-2 ranged from 0.04 to 0.29 (**Table 1**). On average, prediction accuracy was two-fold higher in TBP-1 (0.33±0.08) than TBP-2 (0.16±0.02). For the selected sets (GS, PS and RS), prediction accuracy ranged from 0.25 to 0.32 in TBP-1, whereas it ranged from 0.17-0.29 in TBP-2 (**Table 1**).

### Comparison of phenotypic, genomic, and random selection sets

#### Mean population performance

Across varying levels of proportion selected in TMP, GS (4,658 kg ha^−1^) had significantly higher mean yield than RS (4,548 kg ha^−1^), but none of the differences were statistically significant from PS (4,645 kg ha^−1^) (**Figure 2; Table 2**). The highest mean yield was achieved when selection was stringent (e.g., at 10% selected proportion) at 4,709 kg ha^−1^ and 4,676 kg ha^−1^ for GS and PS, respectively (**Figure 2A; Table 2**). The RS set had consistently lowest mean yield (4,548 kg ha^−1^) regardless of size of selected proportion. In both TBP-1 and TBP-2, there were no significant differences for mean seed yield of GS, PS and RS (**Figure 2A, 2B; Table 2**). Although none of the differences were statistically significant, GS had highest mean seed yield in 2015 and 2015-2016 that differed by at least 6 kg ha^−1^. Based on the 2015 TBP-1 phenotypic data, mean yield of GS (4,983 kg ha^−1^) was higher by 48 kg ha^−1^, 119 kg ha^−1^, and 132 kg ha^−1^ than PS (4,935 kg ha^−1^), RS (4,864 kg ha^−1^) and All (4,851 kg ha^−1^) sets, respectively (**Figure 2; Table 2)**. Similarly, mean yield of GS (5,217 kg ha^−1^) based on the 2015 TBP-2 data, was higher by 6 kg ha^−1^, 16 kg ha^−1^ and 25 kg ha^−1^ than PS (5,211 kg ha^−1^), RS (5,201 kg ha^−1^) and All (5,192 kg ha^−1^) sets, respectively (**Figure 2C; Table 2**). PS was the winning method in 2016 for TBP-1, and differed with respect to GS by 44 kg ha^−1^. Using the two-year data (2015-2016) in TBP-1 and TBP-2, GS had highest mean seed yield over PS and RS, although none of the differences were significant (**Table 2)**. It is worth noting that TBP-2-RS had the highest mean seed yield over other methods for 2016, and nearly identical performance (1 kg ha^−1^ difference) with GS with the 2015-2016 combined data.

**Figure 2.**
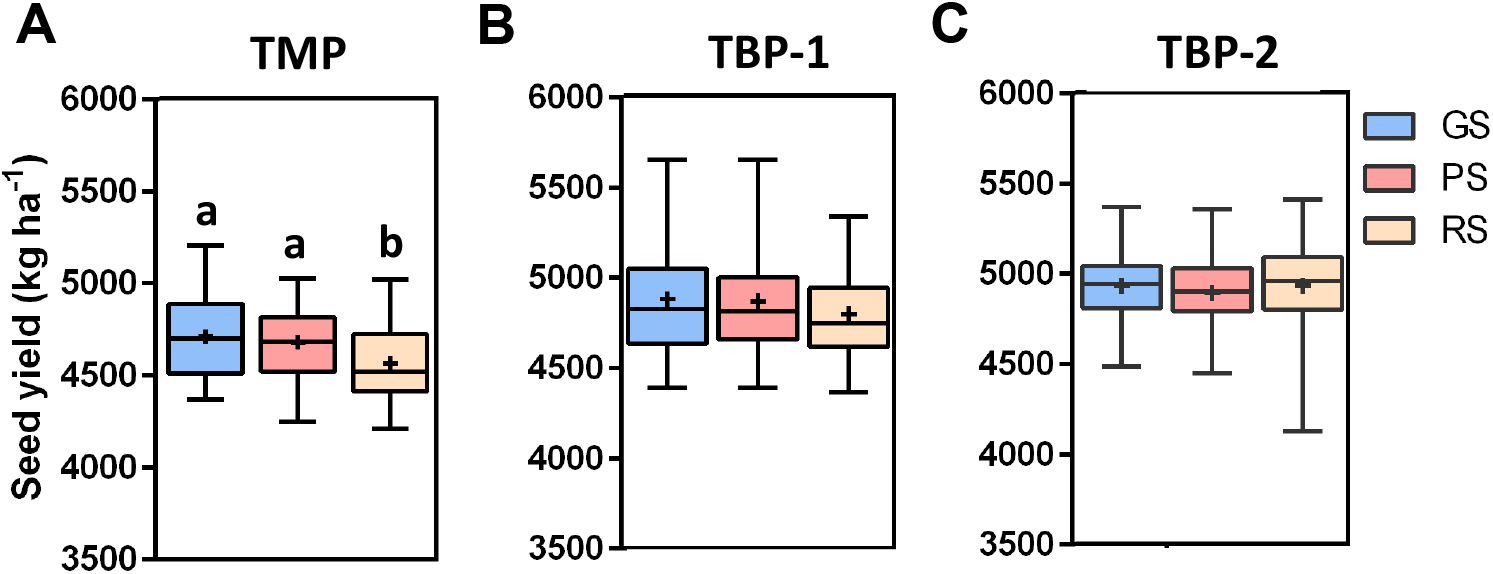
Comparison of seed yield for the selected set at 10% proportion by genomic selection (GS), phenotypic selection (PS), and random selection (RS) in A) TMP, B) TBP-1, and C) TBP-2. TMP was based on the 2013 data with four environments; TBP-1 and TBP-2 were based on 2015-2016 data with 8 environments. Different letters indicate statistically significant differences in mean seed yield (marked by a cross) based on Tukey’s honest significant difference test (a = 0.05). Data within the 25th and 75th percentiles are denoted by the boxes, with line across boxes denoting median and the two tail caps denoting the 10th and 90th percentiles.

**Table 2.**
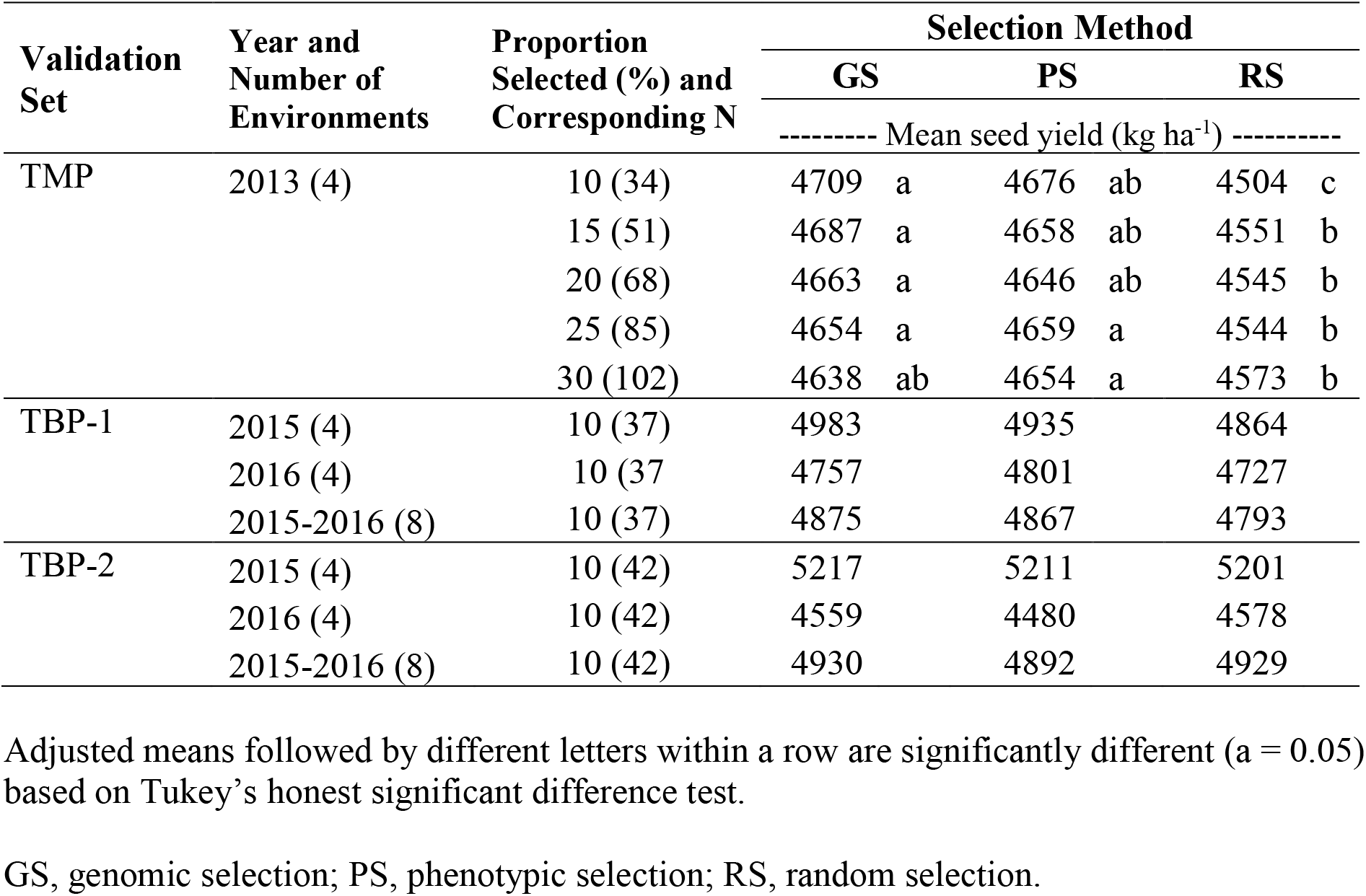
Mean seed yield of the selected sets at different levels of selected proportion in the three validation sets: GS, genomic selection; PS, phenotypic selection; RS, random selection; and SC, all selection candidates tested in the same year. Number of breeding lines (N) is indicated in each selected set.

| Validation Set | Year and Number of Environments | Proportion Selected (%) and Corresponding N | Selection Method |  |  |  |  |  |
| --- | --- | --- | --- | --- | --- | --- | --- | --- |
|  |  |  | GS |  | PS |  | RS |  |
|  |  |  | ----- Mean seed yield (kg ha<sup>-1</sup>) ----- |  |  |  |  |  |
| TMP | 2013 (4) | 10 (34) | 4709 | a | 4676 | ab | 4504 | c |
|  |  | 15 (51) | 4687 | a | 4658 | ab | 4551 | b |
|  |  | 20 (68) | 4663 | a | 4646 | ab | 4545 | b |
|  |  | 25 (85) | 4654 | a | 4659 | a | 4544 | b |
|  |  | 30 (102) | 4638 | ab | 4654 | a | 4573 | b |
| TBP-1 | 2015 (4) | 10 (37) | 4983 |  | 4935 |  | 4864 |  |
|  | 2016 (4) | 10 (37) | 4757 |  | 4801 |  | 4727 |  |
|  | 2015-2016 (8) | 10 (37) | 4875 |  | 4867 |  | 4793 |  |
| TBP-2 | 2015 (4) | 10 (42) | 5217 |  | 5211 |  | 5201 |  |
|  | 2016 (4) | 10 (42) | 4559 |  | 4480 |  | 4578 |  |
|  | 2015-2016 (8) | 10 (42) | 4930 |  | 4892 |  | 4929 |  |

Adjusted means followed by different letters within a row are significantly different ( $\alpha = 0.05$ ) based on Tukey's honest significant difference test.

#### Ranking of best entries

We looked on distribution and frequency of top-yielding lines selected by each method. Frequency of selecting good lines was always higher for GS than PS and RS in all validation sets, except in two out of 11 cases (**Table 3**) where PS was the winning method. In four other cases the difference between GS and PS was only one line. In TMP, the gap between GS and PS in frequency of top performing lines was higher when selected proportion was stringent (e.g., 10-20%). When selected size was 30% of TMP, GS could pick about 42% of top-yielding lines, whereas PS and RS could pick about 38% and 22%, respectively. In TBP-1 and TBP-2, GS had higher frequency of top performing lines in virtually all cases except when selected proportion for TBP-1 was N=30 (**Table 3)**. At best, GS could retain 60% and 40% of top performing lines in TBP-1 and TBP-2, respectively, whereas PS could pick 50% and 30% of top performing lines in TBP-1 and TBP-2 (**Table 3)**.

**Table 3.** Frequency of top-yielding lines that were included in the genomic selection (GS), phenotypic selection (PS), and random selection (RS) sets. Enclosed in parenthesis is the number of tested environments within tested year.

| Validation Set | Year and Number of Environments | Selected Number of Top-Performing Breeding Lines | Selection Method |  |  |
| --- | --- | --- | --- | --- | --- |
|  |  |  | GS | PS | RS |
|  |  |  | Number of lines included in the top-performing breeding lines |  |  |
| TMP | 2013 (4) | 34 | 9 | 4 | 2 |
|  |  | 51 | 15 | 9 | 5 |
|  |  | 68 | 23 | 17 | 9 |
|  |  | 85 | 29 | 28 | 16 |
|  |  | 102 | 43 | 39 | 23 |
| TBP-1 | 2015-2016 (8) | 10 | 6 | 5 | 4 |
|  |  | 20 | 11 | 9 | 6 |
|  |  | 30 | 13 | 14 | 9 |
| TBP-2 | 2015-2016 (8) | 10 | 4 | 3 | 2 |
|  |  | 20 | 6 | 6 | 6 |
|  |  | 30 | 10 | 9 | 8 |

#### Realized cycle gains and relative efficiency

In TMP, realized gain in seed yield for GS and PS was 78 kg ha^−1^ and 74 kg ha^−1^, respectively (**Figure 2A; Table 4**). With varying size of proportion selected in TMP, realized gain was higher when selection was stringent. Highest realized gain in TMP was at 10% selected proportion with realized gain of 129 kg ha^−1^ and 97 kg ha^−1^ in GS and PS, respectively. At 30% selected proportion, PS had higher gain (75 kg ha^−1^) over GS (58 kg ha^−1^). There were 3 cases out of 5 when GS had relative efficiency (RE) greater than 1 over PS in the TMP (**Table 4**). There was no realized gain reported in both TBP-1 and TBP-2 because mean seed yield was not significantly different among methods. However, GS had, for the most part, higher mean seed yield in TBP-1 and TBP-2.

**Table 4.** Realized cycle gain in seed yield in the TMP validation set for phenotypic selection (PS), genomic selection (GS), and random selection (RS) based on four environments.

| Proportion<br>Selected | N | Realized Genetic Gain<sup>‡</sup> |  |  |
| --- | --- | --- | --- | --- |
| | | $GS_R$ | $PS_R$ | $RE_R = GS_R/PS_R$<sup>‡</sup> |
| 10 | 34 | 129 | 97 | 1.3 |
| 15 | 51 | 108 | 78 | 1.4 |
| 20 | 68 | 83 | 67 | 1.2 |
| 25 | 85 | 75 | 79 | 0.9 |
| 30 | 102 | 58 | 75 | 0.8 |

<sup>†</sup> Realized gain was calculated by subtracting the mean of selected set (e.g., GS) from the mean of All set evaluated at year $t$ ;
Percentage gain was calculated as realized gain divided by the mean of All set at year $t$ .
<sup>‡</sup> $RE_R$ = ratio of realized gain by genomic selection (GS) over realized gain by phenotypic selection (PS) at year $t$ ;

#### Genetic similarity

Without any selection, the mean genetic similarity in TMP, TBP-1 and TBP-2 was 0.534, 0.556, and 0.589, respectively (**Table 5**). The observed mean genetic similarity in TBP-1 and TBP-2 was 11% and 18% higher than the expected value of 0.50 in a bi-parental cross. GS had significantly higher mean genetic similarity (TMP-GS at 10%=0.568; TBP-2-GS=0.594) over PS and RS in all validation sets (**Table 5**). On average, GS had at least 2% significant increment in the mean genetic similarity over PS and RS (**Table 5**), suggesting that GS can exhaust genetic variance faster. On the other hand, genetic similarity of TMP-PS selected set (0.532) did not differ significantly from TMP-RS selected set (0.535) at 10% selected proportion. Genetic similarity in the TBP-PS set (TBP-1=0.560; TBP-2=0.586) did not differ significantly from before selection (TBP-1=0.556; TBP-2=0.589) (**Table 5**), indicating that PS may have a minimal loss of genetic diversity. Genetic similarity was significantly lower for RS in TMP (0.523) and TBP-1 (0.547) (**Table 5**), and not significantly different in two other instances.

**Table 5.**
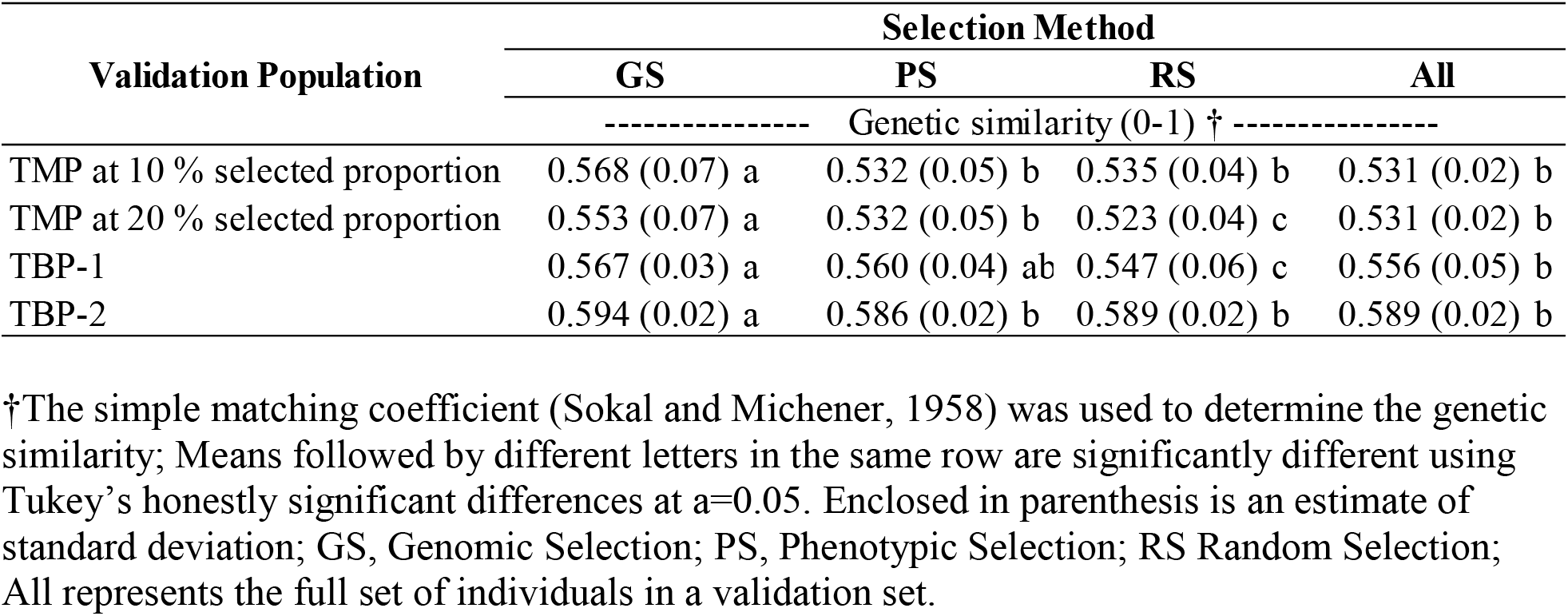
Genetic similarity among individuals included in the selected set of genomic selection (GS), phenotypic selection (PS), random selection (RS) and all individuals included in each selection candidate.

#### Distribution and shift of allele frequency spectrum after selection

Before selection, mean allele frequency of major allele across all SNPs was 0.874 for TMP (**Fig. 3A)** and 0.90 for TBP-1 (**Fig. S3A**) and TBP-2 (**Fig. S3B**). After one round of selection in TMP, GS (0.913), PS (0.885) and RS (0.886) had significantly higher mean allele frequency compared to before selection (0.874) (**Fig. 3A)**. Notably, TMP-GS had significantly higher mean allele frequency than PS and RS. Across all loci, an average increase of 0.04 in allele frequency in the GS set was observed after selection, which is about 5% higher than before selection. The density pattern of allele frequency distribution indicated a significant shift of allele frequency in the GS set towards the extreme (**Fig. 3B,** bottom plot), with more SNP approaching allele fixation (**Fig. 3B,** upper plot). Based on a two-sample, two-sided Kolmogorov-Smirnov (KS) test (*a* = 0.05), we showed that allele frequency distribution was significantly different before (All) and after selection (p<2.2e-16). GS (p<2.2e-16) had the highest upward shift in allele frequency spectrum compared with PS and RS. In general, mean allele frequency values for PS and RS were not significantly different. It is worth noting that RS, as expected, had the smallest density peak (**Fig. 3B**) suggesting less of a shift towards the extreme, compared to other methods, and even less of a shift compared to the set before selection (All). Consistent with what we observed in TMP, mean allele frequency after selection (GS, PS, and RS sets) was higher before selection in both TBP-1 and TBP-2, but we there were no significant differences on mean allele frequency among methods (data not shown).

**Figure 3.**
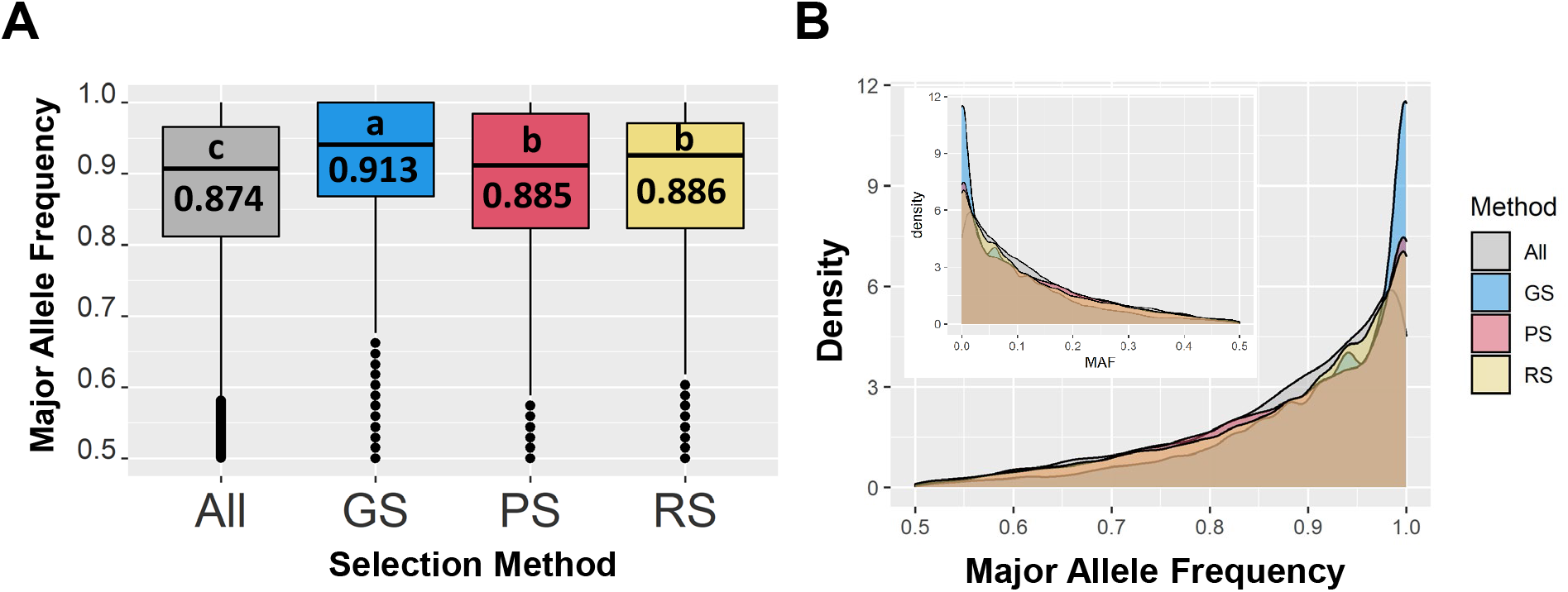
Distribution of allelic frequency in the TMP before (All) and after selection (genomic selection, GS; phenotypic selection, PS; and random selection, RS). (A) Different letters inside each box plot indicate statistical significant differences in mean seed yield based on Tukey’s honest significant difference test (a = 0.05). (B) The bottom density plot represents major allele frequency distribution whereas the upper density plot represents minor allele frequency distribution. Data within the 25th and 75th percentiles are denoted by the boxes, with line across boxes denoting mean, and the two tail caps denoting the 10th and 90th percentiles.

## Discussion

### Prediction accuracy from empirical validation experiments

In this study, we performed a follow-up empirical validation using three independent populations, including a population comprised of pre-selected advanced lines derived from multiple bi-parental crosses, and two independent bi-parental populations. This type of validation experiment requires a significant amount of resources to complete, including the time required for a full breeding cycle. An empirical validation experiment is thus more complex and more thorough compared to typical cross-validation type of experiments, which are abundant in the literature. We found that maximum prediction accuracy (0.54) from our validation experiments was 16% lower than what we obtained from our cross-validation experiments (0.64). Prediction accuracy was consistently higher in TMP and TBP-1, but consistently lower in TBP-2. This is mainly due to the fact that TBP-2 is less related to the TP than TBP-1 and TMP, which is evident in the population structure analysis using PCA (**Figure S2**). In addition, both parents of TBP-1 were extracted from the TP, while one of the parents of TBP-2 was outside the TP. This finding highlights the need for empirical validation experiments, as opposed to simple cross-validation procedures.

### Genomic selection outperforms phenotypic selection at high selection stringency

The goal of selection is to improve the population mean and identify selection candidates with the best genotypic value either through increasing the frequency of favorable alleles (Bernardo, 2009) and/or removal of deleterious mutations (Valluru et al., 2019). Both GS and PS were effective for increasing the population mean for seed yield in TMP, but TMP-GS was significantly different than the population mean before selection in virtually all selection proportions explored. The difference between GS and PS was more pronounced when selected proportion was stringent, suggesting that probability of selecting superior lines using GS was higher when selection intensity was stringent. We did find that frequency of selecting good lines was maximized at 10-20% selected proportion, where TMP-GS could pick a higher number of top-performing lines over TMP-PS. On average, similar realized gains were achieved with TMP-GS (78 kg ha^−1^) and TMP-PS (74 kg ha^−1^).

While we did not see a significant improvement of population mean in both TBP-1 and TBP-2, the biparental TBP-1-GS was actually comparable in gains at 10% selection intensity with the multi-family, more diverse TMP-GS (132 and 129 kg ha-1 for TBP-1-GS and TMP-GS, respectively), despite having a relatively narrower genetic base (Genetic Variance Component of 7.7 and 19.7 for TBP-1-GS and TMP-GS, respectively). This finding provides further support to the effectiveness of GS when using stringent selection intensity. In contrast, gains in the bi-parental population with the highest mean, TBP-2, were minimal with GS and RS and non-existent with PS. For the GS method, the lack of genetic relatedness between the TP and TBP-2 presumably the cause of lower prediction accuracy. The PS method in TBP-2 also resulted an unexpected underperformance with zero realized gains in multi-year evaluations. The observed low performance may be partly attributed to the relatively narrow genetic base of TBP-2, a scenario where genetic improvement tends to be challenging, particularly if the reference population already starts with a high mean, as it is the case. These findings highlight the classical breeding principles of the idealized population, where a breeder may desire to start with a high population mean and also with a diverse population, which in practice is seldom possible.

### Genomic selection leads to greater genetic similarity than phenotypic and random selection

The positive realized gains observed with GS in two of the three independent populations suggest that the observed increase in genetic similarity occurs concomitantly with genetic improvement, and that gains are maximized at higher GS stringency. Stringent GS over cycles could lead to genetic gains and population inbreeding until genetic diversity is exhausted. Our results were consistent with the results obtained in wheat (Rutkoski et al 2015), maize (Jacobson et al 2015), and barley (Sallam et al., 2016) where GS led to a loss of genetic variance. The loss of genetic variance may be due to a significant shift on allele frequency spectrum after selection and leads towards a more rapid allele fixation. A shift on allelic frequency is presumably due to greater linkage disequilibrium between causal variants with effects in the same direction (Brown et al., 2011), which is expected to be prevalent among polymorphism found in bi-parental crosses or closely related individuals. Another possible reason is random genetic drift (Jannink 2010; Rutkoski et al., 2015). GS based on SNP markers may not work on alleles that are not in linkage disequilibrium with markers in the TP, and thus frequencies of such alleles affected by drift (Jannink 2010; Rutkoski et al., 2015). Albeit based on a stochastic simulation, Jannink (2010) observed the same trend in barley where GS could lead to significant loss of many favorable QTL alleles, leading to loss of favorable genetic variance. Placing additional weight on low-frequency favorable alleles can potentially mitigate losses in short-term gain (Jannink, 2010). On the contrary, PS may lead to higher genetic variance than GS because it may act on favorable low frequency alleles, thus increasing frequencies of these alleles more rapidly to intermediate level (Casellas and Varona, 2011; Rutkoski et al., 2015). This may be the reason why PS has been an effective method of selection in maintaining diversity, since it may not drive increases in allele frequencies as hard as GS appears to do, thus, in a way carrying more small-effect favorable alleles that allow for longer-term improvement.

### Towards integration of genomic selection into applied public soybean breeding programs

In applied soybean breeding programs, the goal is to identify the best performing lines that can be used as parents or be directly released as cultivars. Assuming equal realized gain between GS and PS, the implementation of GS may be found beneficial for public breeding programs mainly for shortening the breeding cycle. Assuming GS were used to select new parental lines for the next breeding cycle, this can potentially shorten the cycle time of a typical conventional breeding program by two to three years. One way to approach this is to genotype all selection candidates representing all possible families in the preliminary yield testing stage, then perform selection of candidate parents for the next crossing cycle on the basis of their GEBV. Phenotypic selection will likely be ineffective for selection of seed yield in the preliminary yield test (PYT) stage mainly due to limited amount of available seed at this point, and therefore, new breeding lines could only be tested inevitably in limited number environments with limited to no replications at all. Conducting intermatings at the same time as lines are entering the PYT or before can eliminate two years of phenotypic evaluation, and our empirical data show that GS lines are as good as or superior to PS lines.

Our study was designed to capture the varying genetic relatedness of the TP with validation sets. Consistent from results of several previous studies, GS works best in closely related populations, as in the case of TMP and TBP-1. If selection candidates have less or no closely related progenies with the TP, as in the case of TBP-2, prediction accuracy is expected to be lower, and GS will likely not work as well. To maximize the utility of GS for selection of top-yielding lines, the genomic relationship between the training and any test population needs to be optimally designed. GS will likely work better in closely related breeding populations, mainly comprised of advanced breeding materials, most of which are elite materials, and in which every selection candidate has closely related family (e.g., half-sibs) also being evaluated. In addition, using an optimization algorithm can potentially improve prediction accuracy. For example, following the approach used by Yu et al. (2020), we obtained the reliability criterion for all validation sets, and used reliability values to design two contrasting groups (high vs low reliability set) within each validation population. We consistently found the high reliability sets showing significantly higher prediction accuracy than the low reliability sets in all validation set (**Table S3**).

Genomic selection can lead to a significantly greater loss in genetic variance compared with PS. This is expected, as effective selection reduces genetic variance (Falconer and Mackay, 1996). As favorable alleles become fixed in the elite gene pool, the loss of genetic variance is inevitable due to genetic drift and inbreeding. New mutations are an important source of genetic variation for continued improvement of quantitative traits such as seed yield. Likewise, plant breeders can always replace the genetic variance lost due to allele fixation through introgression of new valuable alleles outside the current breeding program. More research may be needed to develop optimal mating strategies at the varying levels of useable genetic diversity available within the current breeding program.

## Acknowledgements

The authors would like to acknowledge funding for this project from the University of Nebraska-Lincoln, and the Nebraska Soybean Board for funding N. Bandillo’s graduate research assistantship and the ongoing soybean breeding program at the University of Nebraska.

## SUPPLEMENTARY FIGURES

**Figure S1.**
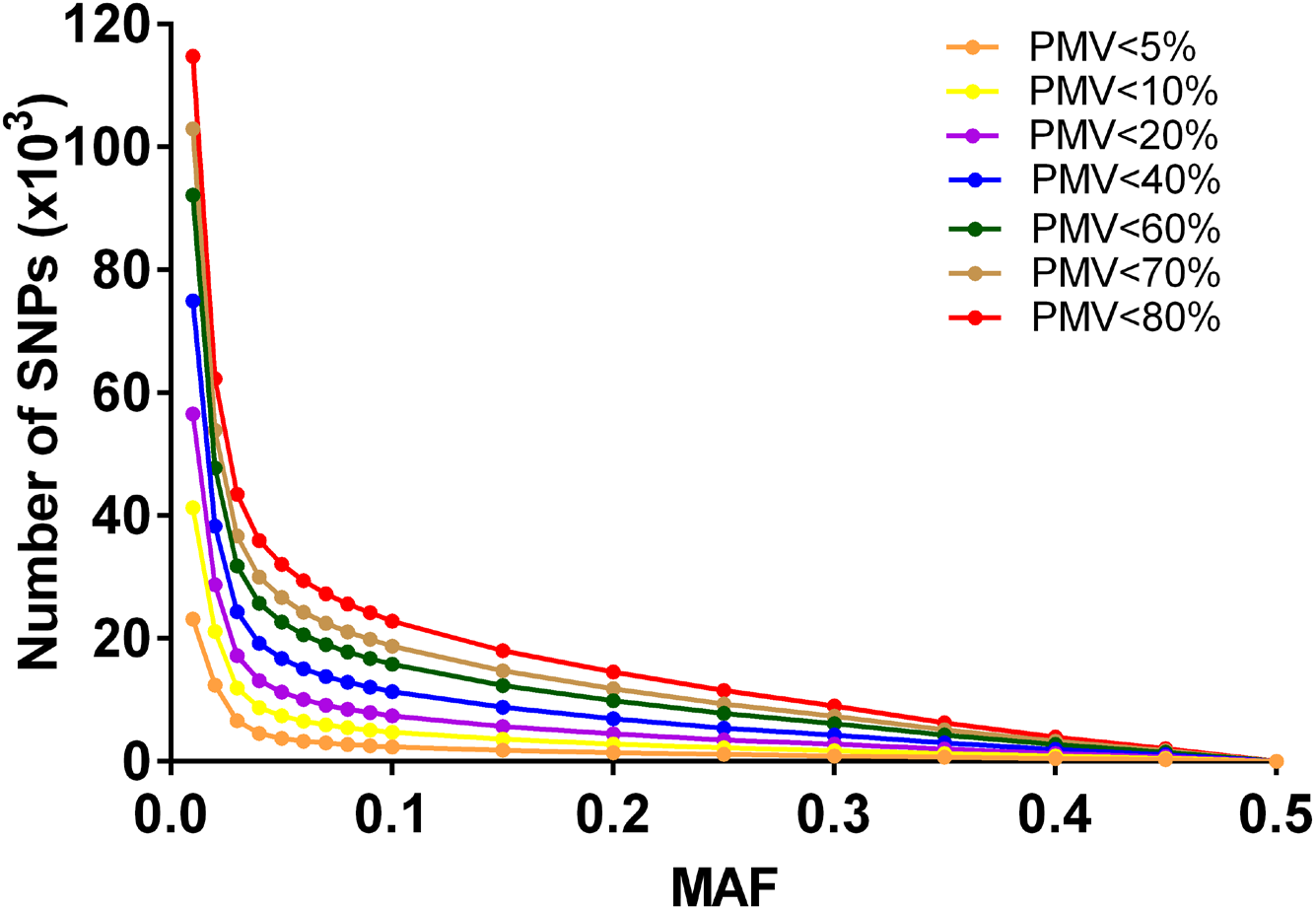
Number of SNPs remaining after applying filtering by combinations of minor-allele frequency and percent missing values

**Fig. S2.**
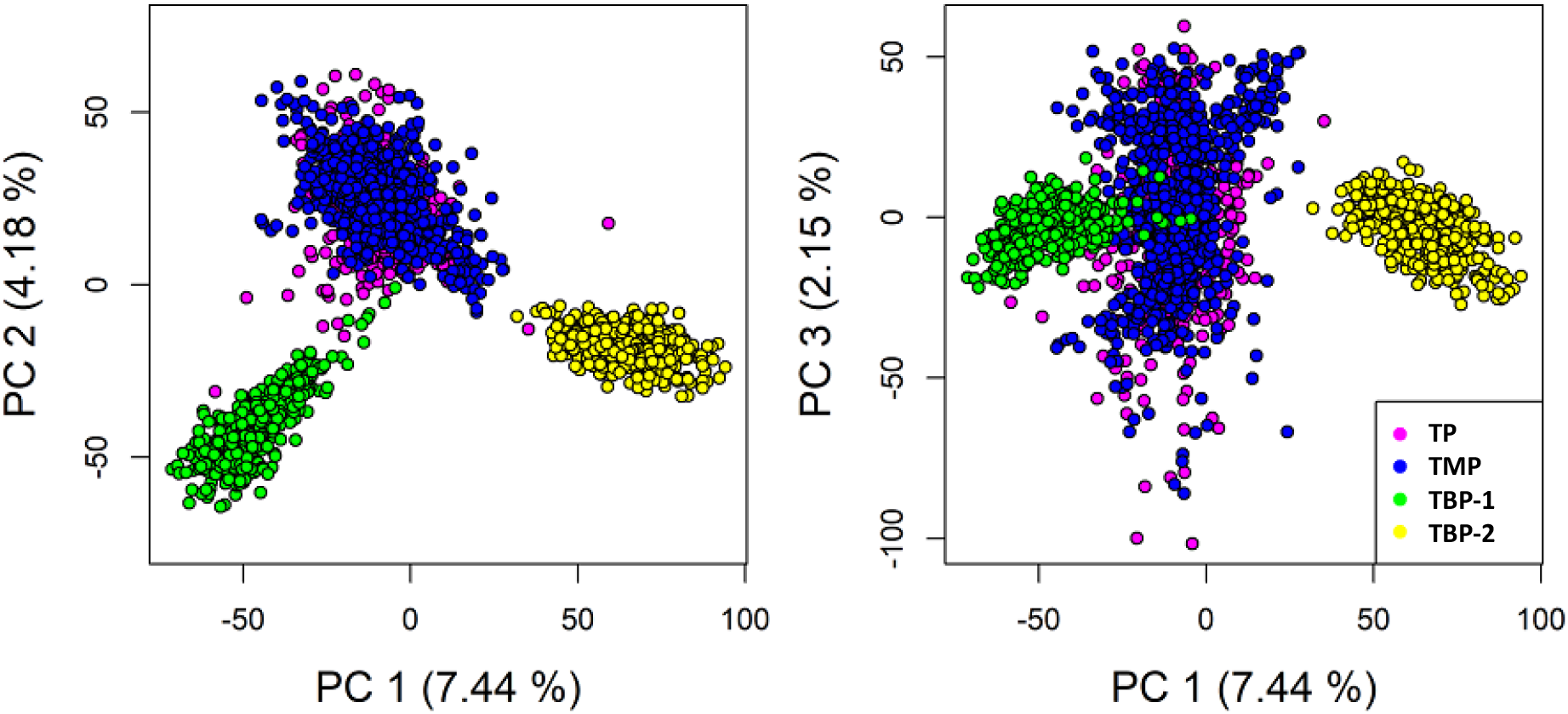
Plant materials and genetic relatedness based on SNP generated from genotyping-by-sequencing.

## SUPPLEMENTARY TABLES

**Table S1.** Set of breeding lines in the training population, validation population, selection candidates, and selected sets by genomic selection (GS), phenotypic selection (PS) and random selection (RS). Population size (N), years of evaluation, experimental design, and number of environments are described in each set.

| Set of Breeding Populations | N | Years of Evaluation | No. of Environments |  |  |
| --- | --- | --- | --- | --- | --- |
|  |  |  | MG 1† | MG 2† | MG 3‡ |
| Training Population | 301 | 2010-2011 | 8 | 8 | 8 |
| TMP | 457 | 2012-2013 | 8 | 8 | 8 |
| TBP-1 (before selection) | 373 | 2014 |  | 4 | 3 |
| TBP-2 (before selection) | 415 | 2014 | 4 | 4 | 3 |
| TBP-1 selected sets | 99§ | 2015-2016 |  | 8 | 8 |
| TBP-2 selected sets | 115§ | 2015-2016 |  | 8 | 8 |

† Lines belonging to maturity groups I and II were evaluated at the Nebraska locations Beemer, Phillips, Cotesfield, and Mead.
‡ Lines belong to maturity group III were evaluated at the Nebraska locations Phillips, Mead, Lincoln, and Clay Center.
§ If a line was selected more than once by at least one selection method, genotypic duplicates were dropped but retained one in subsequent trials.
All environments were laid out following augmented incomplete block design.

**Table S2.** Summary of variance component and heritability for seed yield across different populations in the University of Nebraska soybean breeding program used for genomic selection.

| Breeding Materials | Year | Variance Component ‡ | | | $H^2$ |
| --- | --- | --- | --- | --- | --- |
|  |  | G | GxE | Residual |  |
| Training Population | 2010-2011 | 13.4 | 8.2 | 47.2 | 0.77 |
| TMP before selection | 2012 | 29.8 | 10.1 | 81.3 | 0.70 |
| TMP after selection (GS, PS and RS) | 2013 | 17.6 | 17.1 | 49.0 | 0.63 |
| TMP after selection (GS, PS and RS) | 2012-2013 | 19.7 | 9.8 | 58.5 | 0.80 |
| TBP-1 and TBP-2 before selection | 2014 | 17.5 | 13.8 | 45.5 | 0.66 |
| TBP-1 and TBP-2 after selection (GS, PS and RS) | 2015-2016 | 7.7 | 10.0 | 51.3 | 0.63 |

$H^2$ Broad-sense heritability on an entry-mean basis.
‡G soybean genotype; GxE genotype-by-environment interaction.

**Table S3.** Prediction accuracy (based on G-BLUP) comparison by sampling 100 lines with high reliability and low reliability in the validation population and two independent bi-parental mapping population.

| Validation Set | All | High Reliability Set | Low Reliability Set |
| --- | --- | --- | --- |
|  |  | N=100 | N=100 |
| TMP | 0.41 | 0.49 | 0.26 |
| TBP-1 | 0.43 | 0.45 | 0.41 |
| TBP-2 | 0.13 | 0.34 | -0.02 |

